# Sexual reproduction in the fungal foliar pathogen *Zymoseptoria tritici* is driven by antagonistic density-dependence mechanisms

**DOI:** 10.1101/290072

**Authors:** Frédéric Suffert, Ghislain Delestre, Sandrine Gélisse

**Affiliations:** UMR BIOGER, INRA, AgroParisTech, Université Paris-Saclay, 78850 Thiverval-Grignon, France

**Keywords:** Allocation resource, asexual multiplication, competition, fungal pathogen, plant disease epidemiology, sexual reproduction

## Abstract

This study provides empirical evidence for antagonistic density-dependence mechanisms driving sexual reproduction in the wheat fungal pathogen *Zymoseptoria tritici.* Biparental crosses with 12 increasing inoculum concentrations, in controlled conditions, showed that sexual reproduction in *Z. tritici* was impacted by an Allee effect due to mate limitation and a competition with asexual multiplication for resource allocation. The highest number of ascospores discharged was reached at intermediate inoculum concentrations (from 5.10^4^ conidia.mL^−1^ to 10^6^ conidia.mL^−1^). Consistent with these results for controlled co-inoculation, we found that the intensity of sexual reproduction varied with both cropping period and the vertical position of the host tissues in the field, with a maximum between 25 and 35 cm above the ground. An optimal lesion density (disease severity of 30 to 45%) maximizing offspring (ascospores) number was established, and its eco-evolutionary consequences are considered here. Two ecological mechanisms may be involved: competition for resources between the two modes of reproduction (decrease in the host resources available for sexual reproduction due to their prior use in asexual multiplication), and competitive disequilibrium between the two parental isolates, due to differential interaction dynamics with the host, for example, leading to an imbalance between mating types. A conceptual model based on these results suggest that sexual reproduction plays a key role in the evolution of pathogenicity traits, including virulence and aggressiveness. Ecological knowledge about the determinants of sexual reproduction in *Z. tritici* may, therefore, open up new perspectives for the management of other fungal foliar pathogens with dual modes of reproduction.

## Introduction

An ability to reproduce is one of the most fundamental features of life. It has been suggested that the benefits of sexual reproduction include the purging of deleterious mutations from the genome and the production of a recombinant progeny well-adapted to a changing environment. Density-dependent mechanisms are common in microbial ecology and play a crucial role in sexual reproduction. The modeling of density-dependent selection would shed light on the ability of populations to persist despite “sex”-related conflicts, but simple models seem unable to predict the diversity of the responses observed in nature [1]. A few empirical studies have focused on the role of sex in influencing fungal population and community structures [2, 3].

Several density-dependent mechanisms may operate simultaneously in a population, and it is important to understand how they act, both independently and together. Positive (facilitation) density-dependent mechanisms, such as demographic Allee effects (increasing relationship between overall individual fitness and population density), linked to mate-finding, for example [4], are frequently taken into account in population dynamics studies. Negative (restriction) density-dependent mechanisms, such as competition for resources in cases of high population density, have also been widely described. The effects of competition for resources on individual fitness at low and high densities are a key, complex question in population and evolutionary ecology. For instance, Neiman et al. [5] detected a strong positive effect of high population density on reproductive output in the presence of adequate food supplies in *Potamopyrgus antipodarum* (freshwater snail), but they also identified food limitation as the primary source of negative density dependence. Generally, in situations of high population density, negative density dependence often erases the demographic Allee effect through resource competition.

The impact of density-dependent mechanisms on sexual reproduction has mostly been studied in animal ecology, where encounters between male and female animals result from active behavior influenced by sexual dimorphism or mating advantage [1]. For instance, Stelzer [6] focused on two different mechanisms of density-dependent population regulation — resource exploitation and sexual reproduction — in populations of the rotifer *Brachionus calyciflorus* displaying different investments in sex: populations reproduce clonally at low densities, with the proportion of sexual individuals increasing at higher densities. These two mechanisms have been shown to apply to several facultatively sexual animals, but similar antagonistic density-dependent mechanisms may also operate in fungal microorganisms with dual modes of reproduction.

Competitive interactions and their effect on fitness during reproduction, particularly for asexual reproduction, have been investigated in the emblematic fungal plant pathogens *Puccinia graminis* f. sp. *tritici*, the causal agent of wheat stem rust [7], and *Phytophthora infestans*, the causal agent of potato late blight [8]. The findings obtained were consistent with the ecological principle that resource availability decreases over time, with increasing host tissue mortality, but also during multiple infections, in which different parasite strains share the host resources. Clément et al. [8] focused on the consequences of multiple infections on asexual reproduction of *P. infestans*. By dissociating the effects of single-site and double-site infections and those of competition between identical and different isolates during double-site infections, they showed that the number of strains influences their reproductive rates. Allee effects due to mate limitation have been reported to affect the dynamics of some fungal pathogens at large spatial scales, such as *Hymenoscyphus fraxineus*, the causal agent of ash dieback [9], and *Mycosphaerella fijiensis*, the causal agent of black Sigatoka disease in banana plants [10]. Sexual spores for dispersal are produced only when compatible mating types come into direct contact. However, mate limitation at the plant scale was not confirmed by experimental studies, probably because there are still many gaps in our knowledge concerning the ways in which sexual reproduction occurs in host tissues and its determinants. However, this effect forms the theoretical basis of demogenetic models combining sexual and asexual reproduction, in which the impact of mate-finding limits the speed of pathogen spread [11, 12].

Thus, on the one hand, pathogen population density can affect competition between individuals for limited host resources at a given reproductive stage (e.g. asexual reproduction) or the competition between asexual and sexual stages. On the other hand, it may also affect the probability of finding a mate and, thus, the intensity of sexual reproduction. Antagonistic interactions between negative and positive density-dependent mechanisms have never been studied experimentally in the reproduction of fungal plant pathogens. They could have been investigated by Clément et al. [8] for *P. infestans* co-infections with isolates of mating types A1 and A2 [13], but sexual reproduction was deliberately prevented by inoculation with incompatible isolates only.

Sexual reproduction in *Zymoseptoria tritici*, the causal agent of Septoria leaf blotch of wheat, requires a physical encounter between two compatible strains (Mat1-1 and Mat1-2; [14, 15]) and generally occurs on senescent tissues, with pseudothecia appearing 46 to 76 days after the initial infection in field conditions [16, 17]. It determines the amount of primary inoculum (wind-dispersed ascospores) available for the initiation of subsequent epidemics and plays an essential role in maintaining the diversity of local pathogen populations and determining their rate of adaptation to selection pressures, such as those exerted by resistant hosts or fungicide treatments.

Avirulent *Z. tritici* strains were recently shown to cross with virulent strains on wheat cultivars carrying the corresponding resistance gene [18]. This key finding calls into question current views about the determinants of sexual reproduction, including, in particular, the common belief that individuals unable to infect wheat tissues cannot reproduce sexually. Another recent study has cast doubt on this point given the extent of epiphytic growth seen in some strains, suggesting finally that meeting would be possible without penetration [19]. The proportion of crossing events resulting from the “exclusive paternal parenthood” model [18] was however not quantified at the leaf scale in the natural context of highly diverse populations. In the absence of other factual elements it should be considered that the vast majority of crossing events occur when leaf lesions caused by two compatible strains either coalesce or are located very close together. In practice, it remains unclear how close together infections must be for effective mating to occur. Metcalfe [20] found internal hyphae spanning mean distances of no more than 11 mm in a susceptible cultivar just before the production of pycnidia, suggesting that infection foci located more than 20 mm apart may be isolated. The number of offspring (i.e. the number of ascospores) should be proportional to the number of crossing events, and correlated with disease severity, which is a good proxy for lesion density. This has been shown at field scale: more severe epidemics are associated with higher levels of ascospore production during epidemics in the following year [21, 22]. Moreover, the fitness (aggressiveness) of the *Z. tritici* parental isolates and a time lag between the inoculations with the two parental strains can affect sexual reproduction intensity [23].

The density-dependent mechanisms driving the interaction between *Z. tritici* and wheat during the asexual stage remain unclear. In particular, it remains unknown how the fungus exploits the resources of its host to obtain nutrients for growth, how it invades the apoplast during the long asymptomatic infection phase, and how the necrotrophic phase triggered [24].

### Objective and strategy

We investigated the density dependence of both asexual and sexual reproductive mechanisms in *Z. tritici*, a relevant fungal model organism for such eco-epidemiological investigations. What impact does lesion density in host tissues have on sexual reproduction at the leaf scale? Could we generate empirical data for the establishment of a density-dependence relationship? Would this relationship be monotonic or more complex, due to competition for resources between the two modes of reproduction? This study addressed these questions. We first analyzed epidemiological datasets collected in field conditions, to confirm our initial assumptions. We then assessed the intensities of asexual and sexual reproduction after five *Z. tritici* biparental crosses on adult wheat plants over a period of three years.

## Materials & methods

### Preliminary analysis of field epidemiological data

Field epidemiological data were analyzed to assess the impact of disease severity (proxy of asexual multiplication intensity) and the vertical position of the host tissues on the intensity of subsequent sexual reproduction, during the epidemic period and then during the interepidemic period. We reanalyzed a dataset obtained in Danish field conditions by Eriksen & Munk [16], together with the data obtained in our three-year experiment in France.

*Epidemic period* – The relationship between the proportion of pseudothecia among the total fruiting bodies (pycnidia and pseudothecia) on wheat leaves during a growing season and mean disease severity (expressed as the percentage of the leaf area covered by pycnidia and pseudothecia) were determined with the experimental data for the field experiment performed by Eriksen & Munk [16] (Table 1).

**Table 1.** List of fitness traits and yield components assessed during the asexual and sexual stages for the five *Zymoseptoria tritici* crosses on adult wheat plants.

| Year | Parental isolate | Mating type | Host cv. | Fitness traits |  |  | Yield components |  |
| --- | --- | --- | --- | --- | --- | --- | --- | --- |
|  |  |  |  | Asexual stage |  | Sexual stage | GWS <sup>1</sup> | TKW <sup>2</sup> |
|  |  |  |  | Necrotic area | Density and exudation capacity of pycnidia | Ascospore production |  |  |
| 2015 | INRA09-FS0802<br>INRA09-FS1006 | Mat1-1<br>Mat1-2 | Soissons | × |  | × |  |  |
| 2016 | INRA09-FS0813<br>INRA09-FS0732 | Mat1-1<br>Mat1-2 | Soissons | × | × | × | × | × |
| 2016 | INRA09-FS0808<br>INRA09-FS0806 | Mat1-1<br>Mat1-2 | Soissons | × | × | × | × | × |
| 2017 | INRA09-FS1022<br>INRA09-FS1008 | Mat1-1<br>Mat1-2 | Soissons | × |  | × | × | × |
| 2017 | ST99CH-3D7<br>ST99CH-1A5 | Mat1-1<br>Mat1-2 | Runal | × |  | × | × | × |

<sup>1</sup> grain weight per spike
<sup>2</sup> thousand-kernel weight
<sup>3</sup> also assessed on two sets of plants inoculated separately with each monoparental suspension

*Interepidemic period* – In a field plot sown with a monoculture of wheat cv. Soissons from 2007 to 2016 at the Grignon experimental station [22], standing stubble was left over an area of 20 m^2^ during the fall of 2015-2016, 2016-2017 and 2017-2018. Stubble samples were collected three or four times each year, from September to January. Stems and leaves were partitioned from lower to upper layers, according to their vertical position (0-5 cm, 5-10 cm, 10-15 cm, 15-25 cm, 25-35 cm, 35-50 cm, 50-65 cm). The dry tissues of each layer of wheat stubble were weighed, and ascospore production, as a proxy of sexual reproduction intensity, was quantified as described by Suffert et al. [17].

### Fungal material

Five crosses were performed, with eight *Z. tritici* isolates (FS0802 × FS1006, FS0813 × FS0732, FS0808 × FS0806, FS1022 × FS1008 [25]) collected in 2010 from wheat cv. Soissons in France, and two isolates (3D7 × 1A5 [26]) collected in 1999 from wheat cv. Galaxie and Lono (Table 1). The compatibility of the five pairs of isolates was determined by PCR amplification of the two mating-type idiomorphs [15]. The effective ability of the eight French isolates to reproduce sexually on adult plants was first checked in semicontrolled conditions [23]. Subcultures of each isolate stored at −80°C were grown for five days in Petri dishes containing PDA (potato dextrose agar, 39 g L^−1^) at 18°C in the dark. For each isolate, an initial conidial suspension was prepared with a Malassez counting chamber. Serial dilutions (1:5) of each initial monoparental suspension were prepared, from 10^7^ conidia mL^−1^ to 50 conidia mL^−1^. Twelve biparental suspensions were then prepared by mixing 25 mL each of the appropriate monoparental suspensions and adding two drops of surfactant (Tween 20; Sigma, France).

### Plant material

Seeds of the wheat cv. Soissons or Runal (moderately susceptible to Septoria tritici blotch) were sown on December 18 2014, December 14 2015, and December 15 2016 (Table 1) in Jiffy peat pots, which were then kept in greenhouse conditions for two weeks. Seedlings were vernalized in a growth chamber for 8 weeks at 8°C, with a 10-h light/14-h dark photoperiod. They were then returned to the greenhouse and left to acclimate for one week before transplantation into individual pots filled with 1 liter of commercial compost (Klasmann Peat Substrat 4; Klasmann France SARL, France). We added 4 g of slow-release fertilizer (Osmocote Exact 16-11-11N-P-K 3MgO Te) to each pot. The plants were also watered with Hydrokani C2 fertilizer (Hydro Agri Spécialités, France), diluted 1:100 and poured into the saucers under the pots. Plants were sprayed once with spiroxamine (Flexity SC at 300 g L^−1^; BASF, France) for the specific prevention of powdery mildew *(Blumeria graminis* f.sp. *tritici)*, no later than six weeks before inoculation. During the growth period, the plants were illuminated with natural daylight supplemented with 400 W sodium vapor lamps between 6.00 a.m. and 9.00 p.m. The air temperature was kept below 20°C during the 15-h light period and above 12°C during the 9-h dark period. Plants were thinned to three stems per pot during the growth period.

### Inoculation procedure

Co-inoculations with the five biparental suspensions were carried out after the wheat heads had fully emerged (one in 2015, on April 30; two in 2016, on May 3; two in 2017, on April 25; Table 1). In 2017, two complete sets of wheat plants cv. Soissons were also inoculated with each monoparental suspension (FS1022 and FS1008). The suspensions were applied with an atomizer (Ecospray, VWR, France), on three adult plants (nine stems), as described by Suffert et al. [23]. Plants were turned during the 10-second spraying event, to ensure even coverage with inoculum. Infection was promoted and cross-contamination prevented by enclosing each trio of plants inoculated with the same suspension in a sealed transparent polyethylene bag containing a small amount of distilled water for 72 h. Wheat plants were then kept in a greenhouse for about 13 weeks (from April 30 to July 24 2015, from 3 May to July 28 2016, from 25 April to August 2 2017; Figure 1a). Air temperature was kept above 18°C during the spring, and reached 30°C several times during summer.

**Fig. 1.**
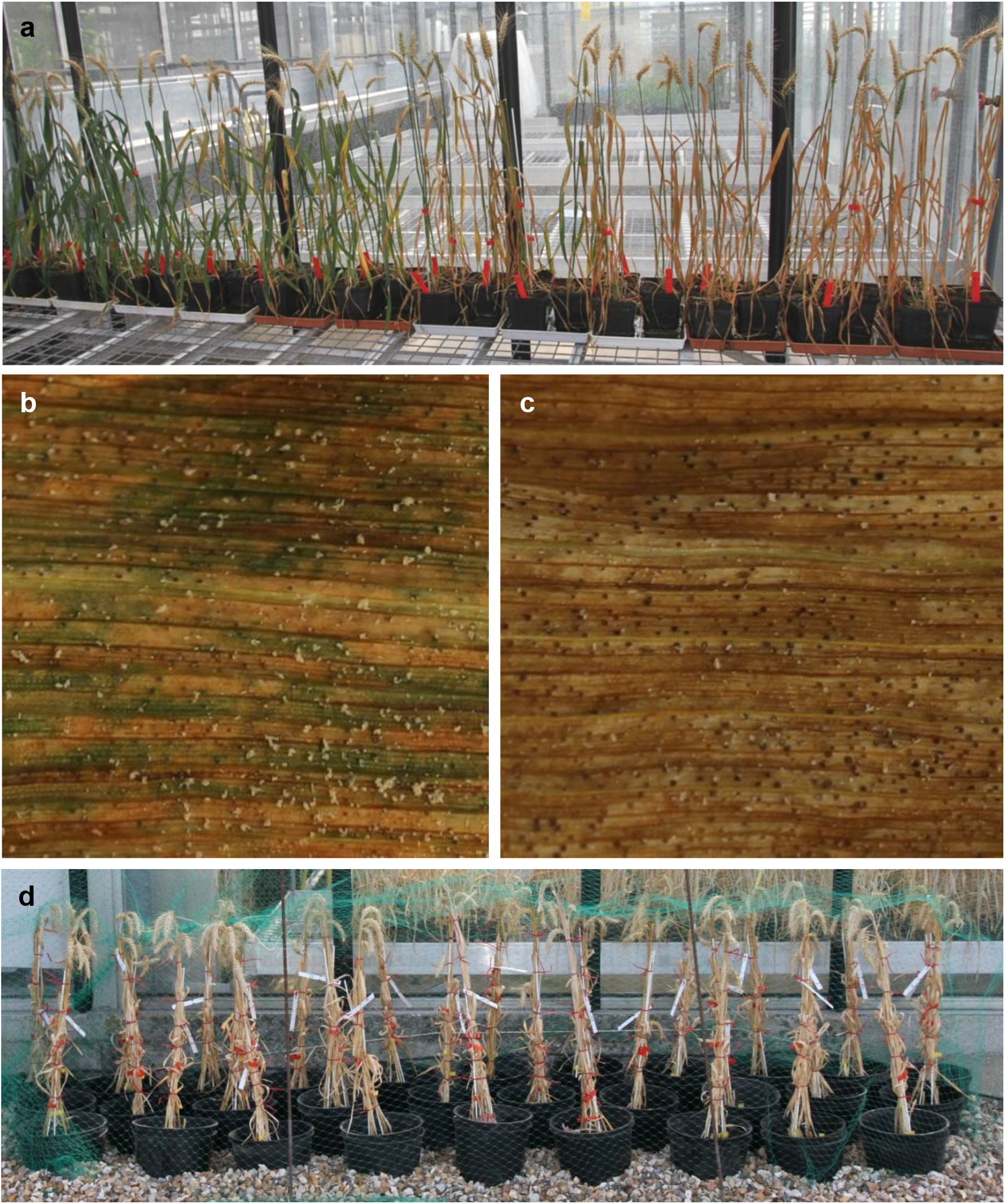
(a) Adult wheat plants cv. Soissons (three replicates per inoculum concentration) five weeks after their inoculation in the greenhouse with 12 biparental suspensions of *Zymoseptoria tritici* isolates (FS808 and FS806), at concentrations of 50 to 10^7^ conidia.mL^−1^. Plants are ranked from the lowest (left) to the highest concentration (right). (b-c) Sporulating area on the flag leaf, characterized by a high density of pycnidia, with high (b) and low (c) exudation capacities. (d) Trios of plants stored outdoors to induce sexual reproduction and ascosporogenesis.

### Assessment of fitness traits during the asexual stage

Disease severity on leaves was assessed by eye five weeks after inoculation (on June 2 2015, June 6 2016 and May 27 2017), as the percentage of the leaf area displaying necrosis (mean for the F1, F2, and F3 leaves for each stem; 1, 2, 3 and 5%, thereafter by increments of 5% up to 100%; Figure 2). In 2016, the density of pycnidia (number of pycnidia divided by the necrotic area) and the proportion of pycnidia exuding a cirrhus (Figure 2b-c) were also estimated for plants inoculated with each of the two biparental suspensions, FS0813 × FS0732 and FS0808 × FS0806 (Table 1), from the necrotic area in a 6-cm leaf section from four randomly selected F1 leaves per inoculum concentration.

**Fig. 2.**
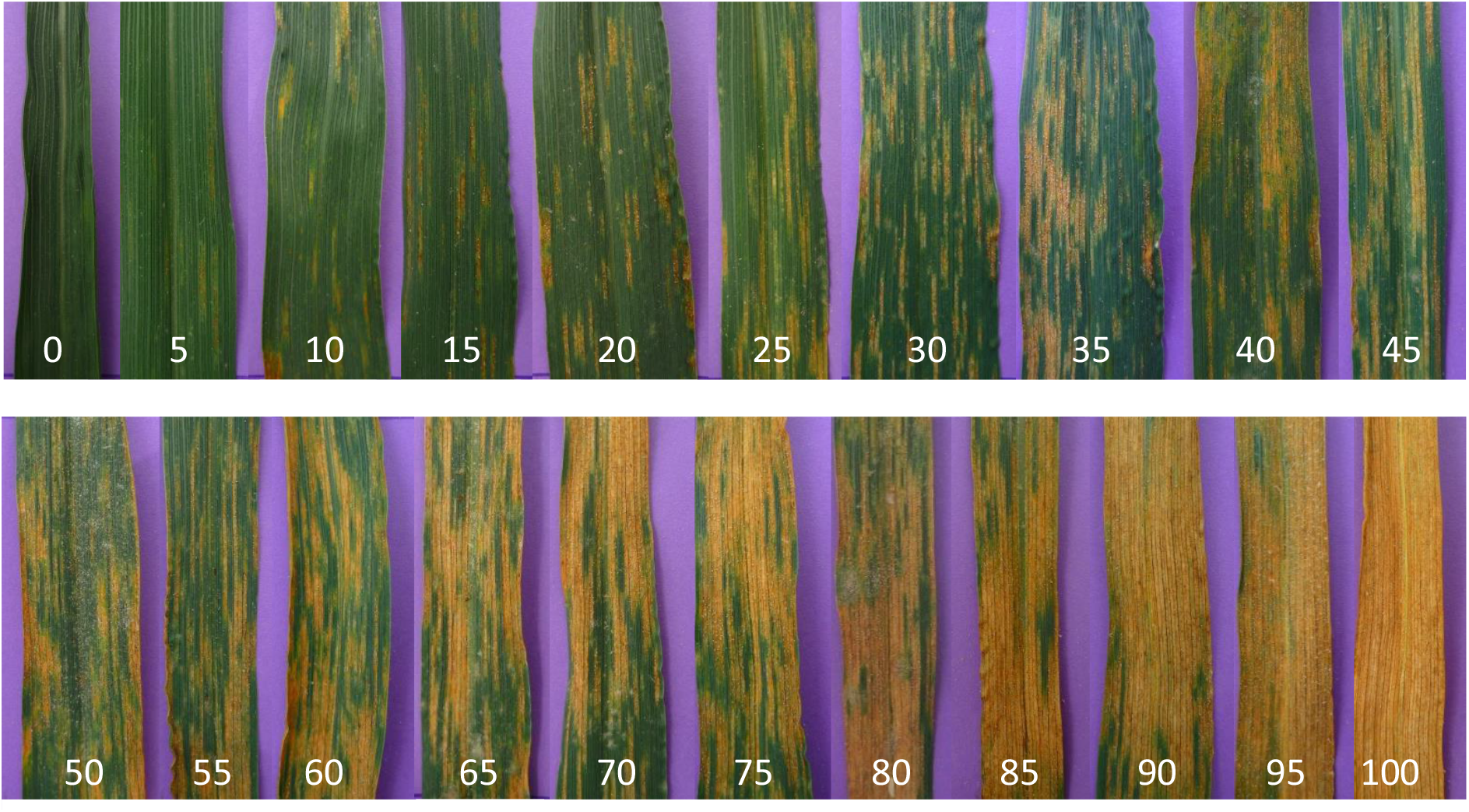
Visual rating scale used to quantify the intensity of *Zymoseptoria tritici* asexual multiplication (disease severity assessed as a percentage of necrotic area) on the three uppermost leaves (F1, F2 and F3) of adult wheat plants. The images show the 2-7 cm section of the flag leaves over their point of insertion on the stem.

### Promotion of sexual reproduction and ascosporogenesis

Co-inoculated plants were placed outdoors during the summer and fall (from July 24 2015 to October 9 2015, from July 28 2016 to November 14 2016, and from July 28 2017 to December 4 2017) to induce the maturation of pseudothecia (Figure 1d). Each trio of coinoculated plants was tied up with stiff wire, with the stems and leaves clamped so as to prevent contact with other trios of plants. On October 14 2015, November 16 2016, and December 6 2017, dry leaves and stems from each trio of plants were cut into 2-cm fragments and allowed to dry, in separate open boxes, in a laboratory atmosphere, at 18°C for one week. Ascospore release was assessed immediately after drying, to check the maturity of the pseudothecia. The pseudothecia were considered mature in 2015 and 2017. By contrast, in 2016, the plant debris was placed outdoors again until January 16 2017, to promote pseudothecium maturation.

### Assessment of the intensity of sexual reproduction

For each inoculum concentration, the debris was weighed, soaked in water for 20 min and spread on dry filter paper in a box (24 × 36 cm), the lid of which was left half open to decrease the relative humidity of the air around the debris. Eight Petri dishes (90 mm in diameter) containing PDA medium were then placed upside down 1 cm above the debris. The boxes were placed in the dark, at 18°C for 19 h. The Petri dishes were then closed and incubated in the same conditions. Each discharge event was replicated four times (from November 2015 to January 2016, from January 2016 to February 2017, and from December 2017 to January 2018).

The ascospores released onto the Petri dishes germinated after 24 h. Six days later, yeast-like colonies resembling cream-colored convoluted heaps were observed. The colonies were counted under a microscope four and seven days after ascospore discharge. It was assumed that each colony resulted from the germination of a single ascospore, and that clusters of colonies appeared just above mature pseudothecia from which several ascospores had been discharged. A cumulative ascospore discharge index (ΣADI), defined as the total number of ascospores discharged per gram of wheat debris [17, 23], was calculated for each discharge event, as follows:

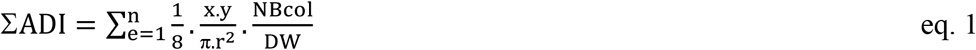

where NBcol is the total number of colonies in the eight Petri dishes, DW is the total dry weight of debris spread out in the box, *r* is the radius of a Petri dish (4.5 cm), x and *y* are the width and length of the box (24 cm and 36 cm, respectively), and *n* is the number of discharge events (*n* = 4).

### Assessment of yield components

Yield components were assessed in 2016 and 2017 (Table 1). At maturity, the spikes were cut and hand-threshed to separate the kernels from the chaff. Kernels were then counted and weighed to estimate the grain weight per spike (GWS) and thousand-kernel weight (TKW) for each cross and inoculum concentration.

### Data analysis and visualization

Statistical analysis - ANOVA or Kruskal-Wallis test when relevant and post hoc multiple comparisons - were performed using the R software.

A quadratic equation (eq. 2) was used to visualize the relationship between sexual reproduction intensity (SRI) and asexual multiplication intensity (AMI).

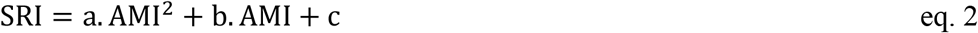

where *a, b* and *c* are parameters.

An alternative equation integrating two types of density-dependent constraints (eq. 3) was also used. It was based on a logistic function f(α,β,γ,AMI) (eq. 4) describing a positive density-dependent mechanism and coupled with a function g(μ,σ,p,AMI) (eq. 5) describing a negative density-dependent mechanism.

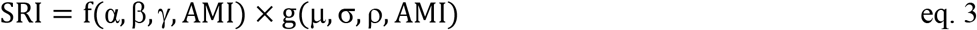

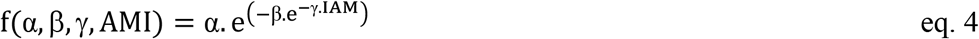

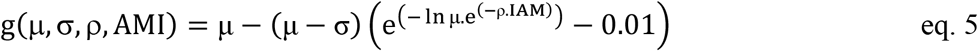

where α, β, γ, μ, σ, and ρ are parameters.

Both equations were adjusted with the data generated by the five crosses, with the estimation of SRI by determination of the cumulative ascospore discharge index (ΣADI) and of AMI with the percentage of the total area displaying necrosis.

## Results

### Sexual reproduction in field conditions varies according to the cropping period and the vertical position of the infected host tissues in the canopy

The epidemiological data obtained by Eriksen & Munk [16] in field conditions showed that the proportion of pseudothecia (proxy for the intensity of sexual reproduction) present on wheat leaves varies with disease severity (proxy for asexual multiplication intensity; Figure 3) over the cropping season. Our reanalysis of these data made it possible to limit bias due to fruiting dynamics (the latent period is much longer for pseudothecia than for pycnidia). The proportion of pseudothecia was maximal for severities of 50 to 80%. This proportion remained stable or even decreased at severities above 80%.

**Fig. 3.**
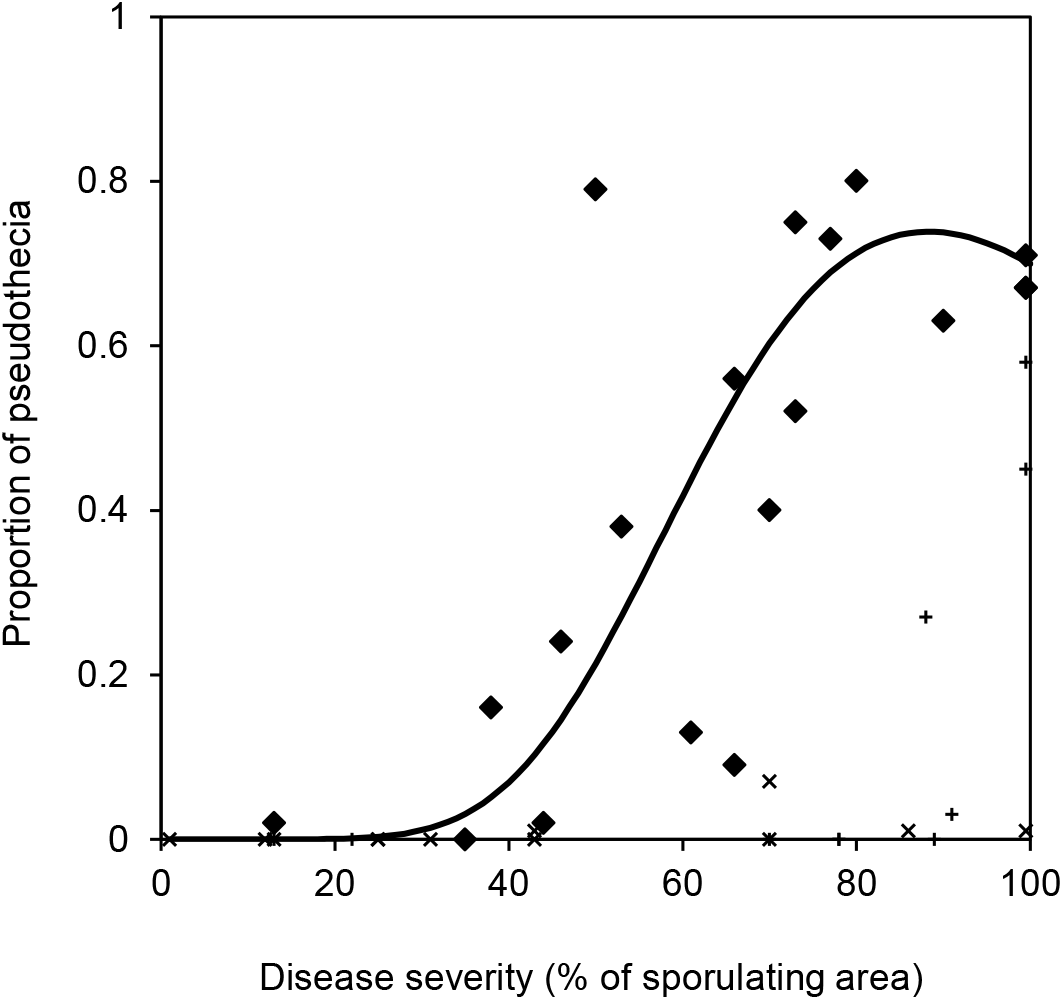
Relationship between the proportion of pseudothecia among the total fruiting bodies on wheat leaves in field conditions and the mean severity of Septoria tritici blotch expressed as a percentage of the leaf area covered by pycnidia or pseudothecia. An equation integrating different density-dependent constraints (bold line; see eq. 3 in Material & Methods) was fitted to some of the experimental data (♦) obtained by Eriksen et al. [16] (Table 1) in a two-year field survey in Denmark. Data for leaves L8 to L11 (basal leaf layers) were excluded (×) because no pseudothecia were found whatever the level of disease severity, probably due to their advanced state of decomposition; data for leaves L5 to L7 (collected before mid-June), L1 (flag leaf), and L2 (collected before August), were also excluded (+), because sexual reproduction was probably incomplete (29-53 days between the appearance of the first pycnidia and the appearance of the first pseudothecia [16]).

Monitoring of the dynamics of sexual reproduction on standing stubble over three interepidemic periods revealed a vertical structuring of mean ascospore production on senescent wheat tissues (Online Resource 1). This vertical structuration changed over time. Ascospore numbers peaked in November (2015, 2016) or December (2017). Ascospore production generally increased with distance from the ground, up to a height of 25 cm (Figure 4). This layer corresponds to the part of the stubble not usually exported at harvest. The 25-35 cm layer made the largest contribution to sexual reproduction. Above this height, ascospore production follows the opposite trend, gradually decreasing between 35 and 65 cm. The lowest layer of standing stubble, which includes the crown and what remains of the first leaves to emerge before tillering, generated less than 5% of the total number of ascospores.

**Fig. 4.**
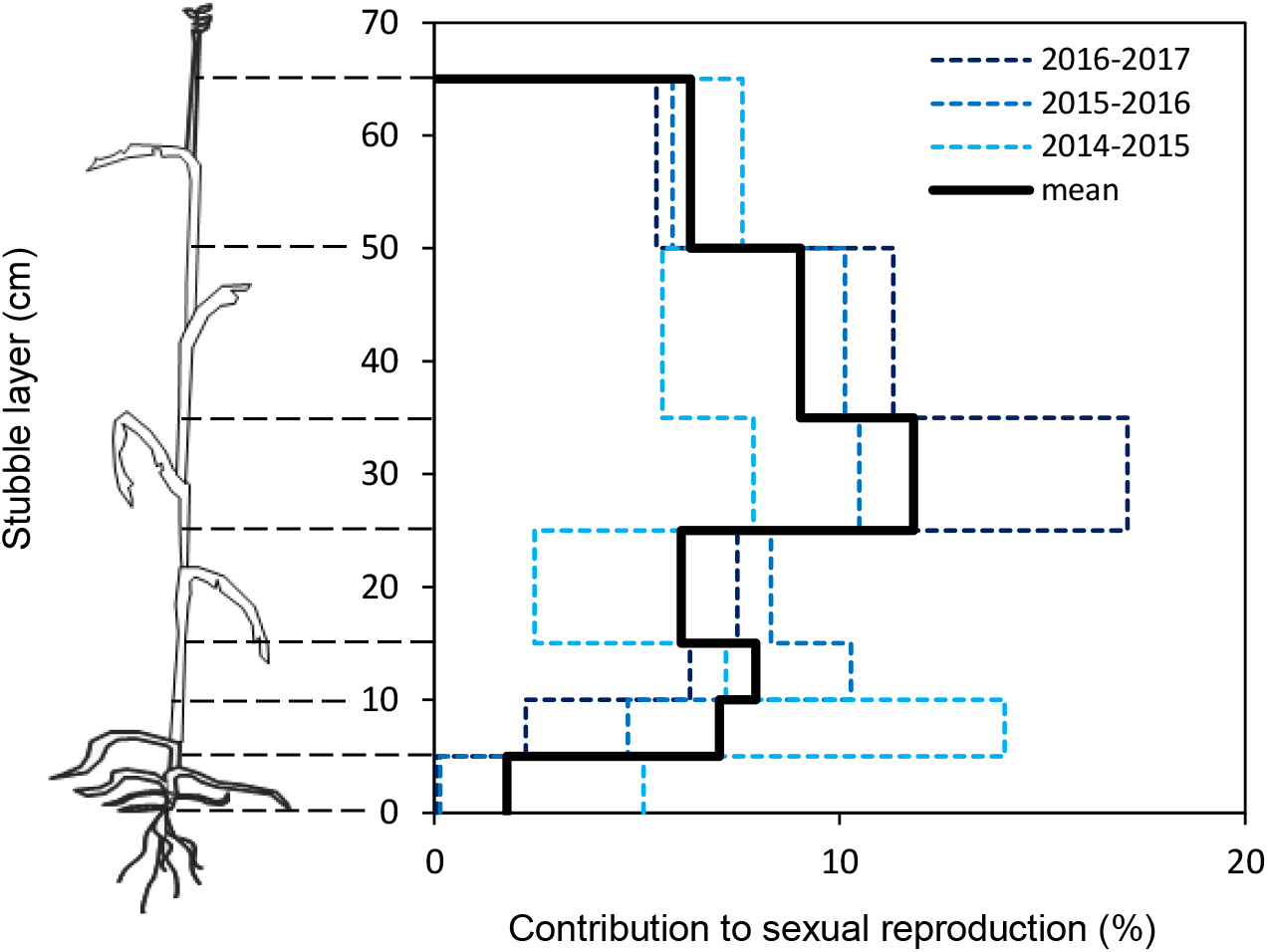
Impact of the vertical position of wheat tissues in standing stubble left in the field in the fall (stems and leaves, from lower to upper layers: 0-5 cm, 5-10 cm, 10-15 cm, 15-25 cm, 2535 cm, 35-50 cm, 50-65 cm) on the intensity of *Zymoseptoria tritici* sexual reproduction expressed as the number of ascospores collected per gram of debris (logarithmic scale) on three or four dates (from September to January) over three interepidemic periods (2015-2016, 2016-2017, 2017-2018). The mean contribution of each stubble layer to sexual reproduction is expressed as a percentage of the total number of ascospores collected during each interepidemic period.

### The general interaction between *Z. tritici* and wheat is density-dependent

The interaction between the pathogen and the host plant is clearly density-dependent. Regardless of the year in which the cross was performed (2015, 2016 or 2017), the biparental suspension used and the host cultivar (Soissons or Runal), inoculum concentration (total number of *Z. tritici* conidia in the biparental suspension per mL) was positively related to the estimated mean disease severity (percentage of necrotic area) for the three uppermost leaves of the adult wheat plants (Figure 5). Soissons was particularly susceptible to the isolates with which it was co-inoculated (mean sporulation area of 60% for an inoculum concentration of 5 × 10^5^ conidia.mL^−1^), whereas Runal was more resistant (35%). For both varieties, disease severity (80%) was maximal at an inoculum concentration of 10^7^ conidia.mL^−1^.

**Fig. 5.**
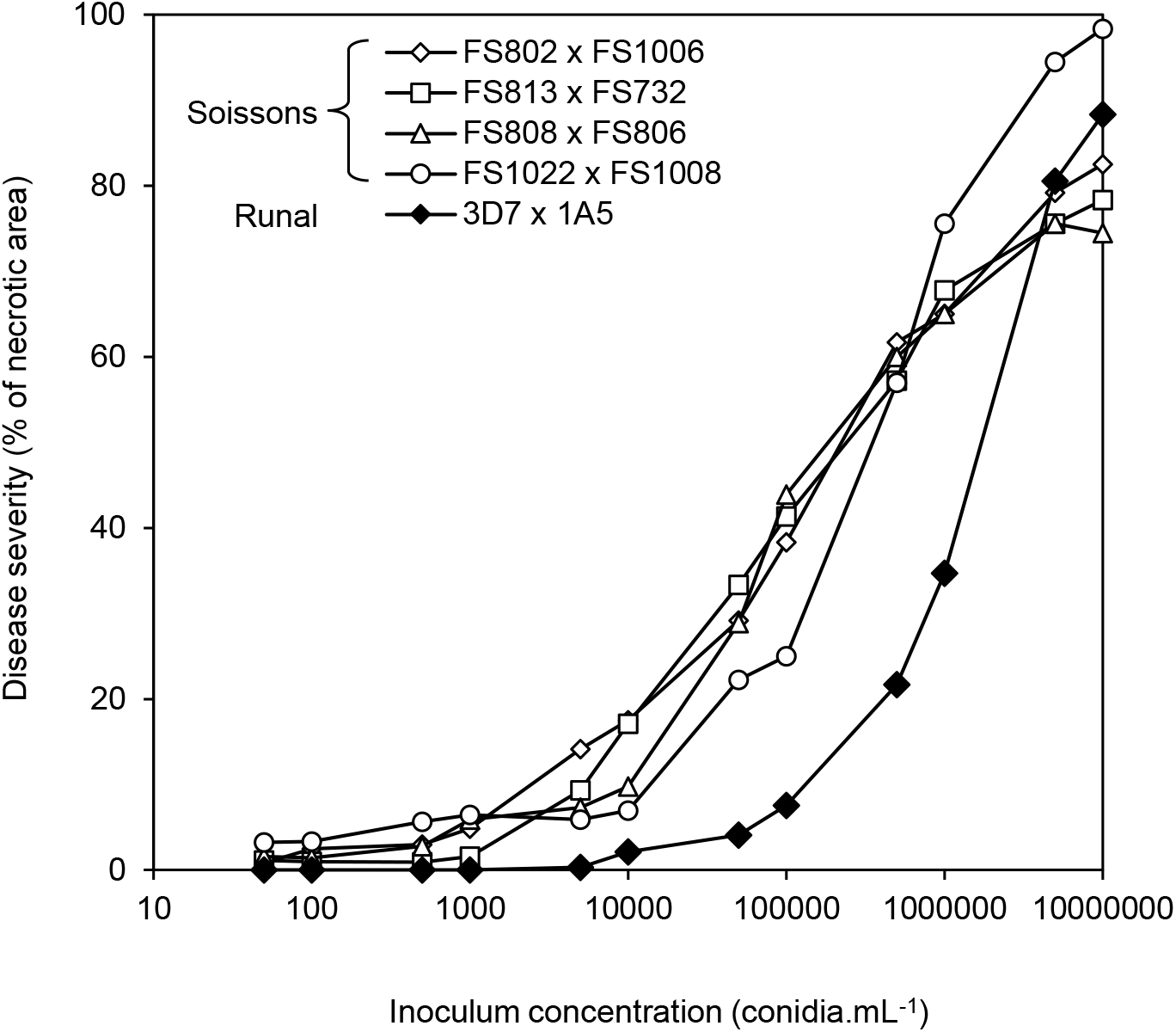
Relationship between *Zymoseptoria tritici* inoculum concentration (total number of conidia per mL) and mean intensity of asexual multiplication (disease severity assessed as a % of necrotic area) estimated on the three uppermost leaves of adult wheat plants. Inoculation with biparental suspensions of isolates FS802 × FS1006 (2015), FS813 × FS732 (2016), FS808 × FS806 (2016), and FS1022 × FS1008 (2017) were performed on cv. Soissons; Inoculations with biparental suspensions of isolates 3D7 × 1A5 (2017) were performed on cv. Runal.

Mean severity on the three uppermost leaves of adult wheat plants cv. Soissons and Runal was inversely correlated with total grain weight per spike (GWS), and thousand kernel weight (TKW): late (inoculation after flowering) and intense attacks (sporulation over 80% of the area) decreased grain yield by up to 20-25% (Figure 6). *Z. tritici* attacks had a strong impact on wheat yield, demonstrating that the interaction between the pathogen and the host plant was density-dependent, i.e. a function of pathogen inoculum pressure.

**Fig. 6.**
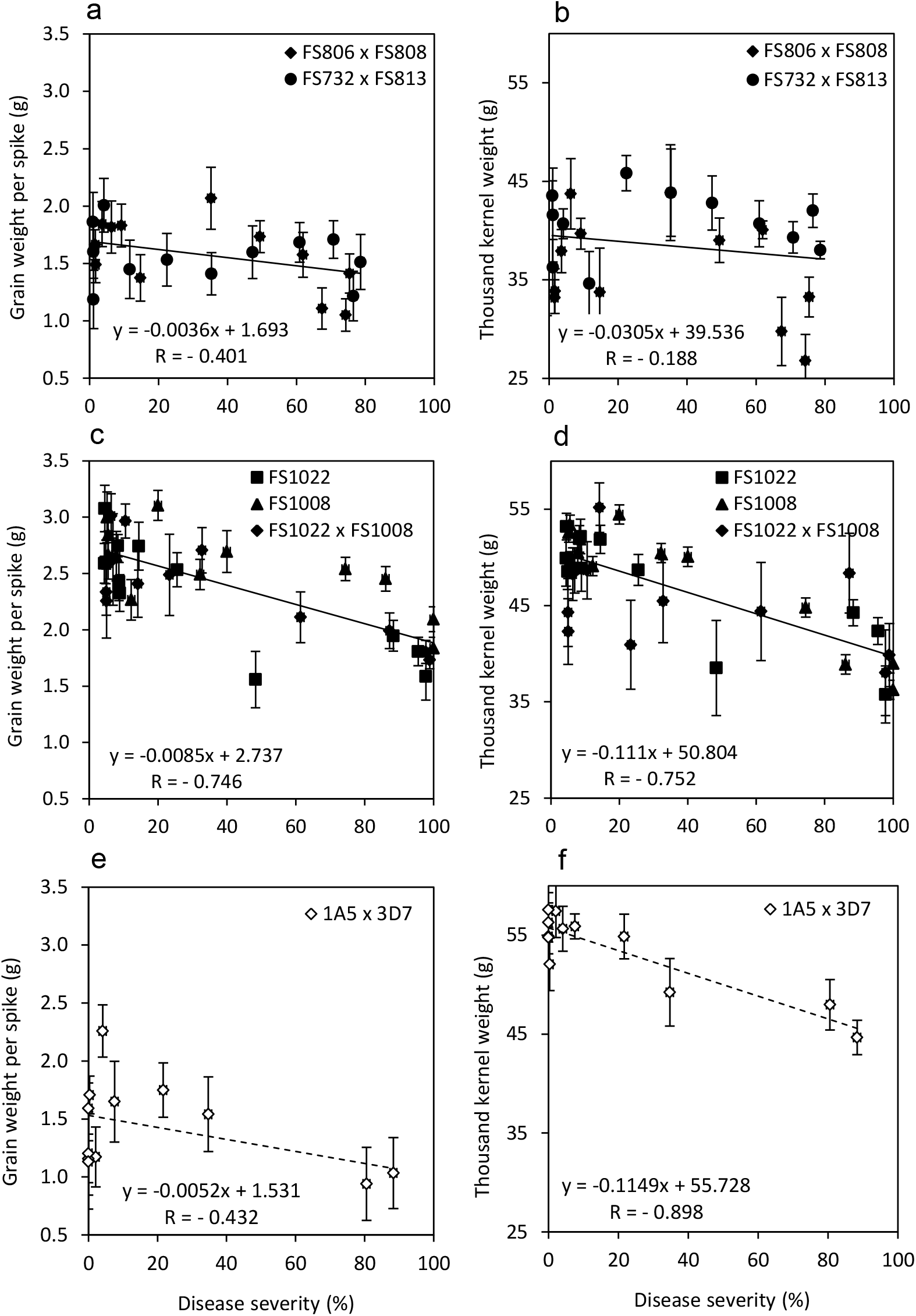
Relationship between the mean intensity of asexual multiplication in *Zymoseptoria tritici* (disease severity assessed as a % of necrotic area) on the three uppermost leaves of adult wheat plants cv. Soissons (black symbols) or Runal (white symbol), and (a, c, e) total grain weight per spike (GWS) or (b, d, f) thousand-kernel weight (TKW). Disease severity was assessed five weeks after inoculation with monoparental or biparental suspensions of isolates FS806, FS808, FS732, and FS813 in 2016, and FS1022, FS1008, 1A5, and 3D7 in 2017. Vertical bars represent the standard deviation. The Pearson correlation coefficient (R) is given for each relationship.

### Competition for resources affects asexual multiplication in *Z. tritici*

We observed robust, negative relationships between mean disease severity and the density of pycnidia within lesion, and between mean disease severity and the percentage of exuding pycnidia (i.e. density of cirrhi within lesion) in 2016 (Figure 7): more intense *Z. tritici* attacks were associated with a lower density of pycnidia, and a lower percentage of exuding pycnidia. For both FS813 × FS732 and FS808 × FS806 crosses, severe attacks (mean sporulation area of 80%) halved the density of pycnidia (150 vs. 300 pycnidia.cm^−2^) and the percentage of exuding pycnidia (45 vs. 90%). Thus, the asexual development of *Z. tritici* is affected by competition for resource allocation.

**Fig. 7.**
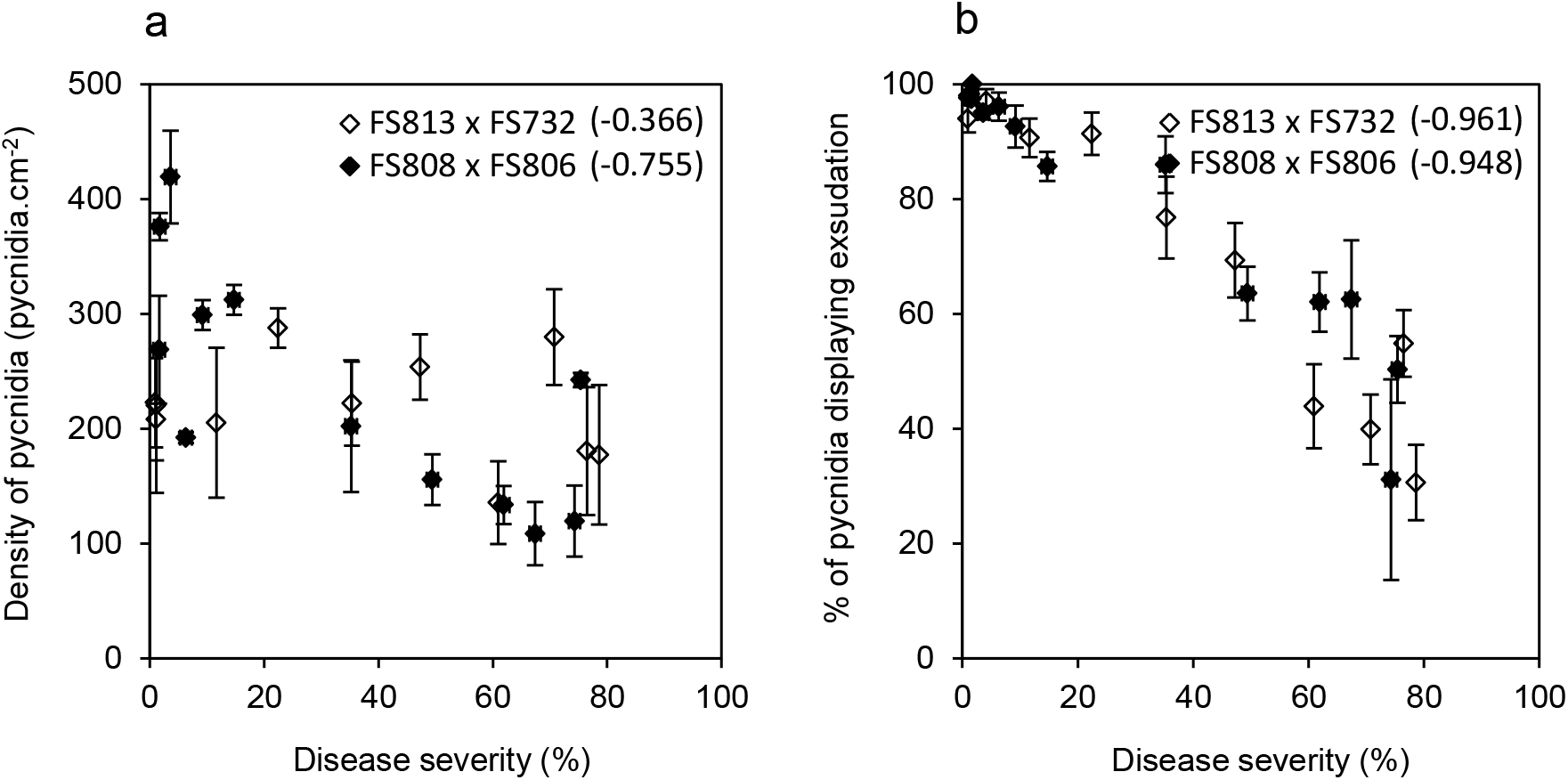
Relationship between the mean severity of Septoria tritici blotch (percentage of necrotic area) and (a) density of pycnidia (number of pycnidia per cm^2^) or (b) percentage of pycnidia displaying exudation. Assessments were performed five weeks after the inoculation of adult wheat plants cv. Soissons with biparental suspensions of *Zymoseptoria tritici* isolates FS813 × FS732 (2016) and FS808 × FS806 (2016). Vertical bars represent the standard deviation. The Pearson correlation coefficient is given in brackets.

Disease severity reached after inoculation with an isolate inoculated alone (FS1008) was compared to the disease severity reached after inoculation with a biparental suspension (FS1008 × FS1022) containing the same number conidia of the previous isolate (FS1008). Each of the five groups of plants that were compared was thus exposed to a single or double *Z. tritici* inoculum pressure, but to the same amount of FS1008 conidia. The comparison highlighted competitive disequilibrium between the two parental isolates (Figure 8). This finding provides a second line of evidence that the asexual development of *Z. tritici* is driven by density-dependent processes. Soissons appeared to be more susceptible to FS1008 than to FS1022, regardless of the concentration of the inoculum: in practice, FS1008 was more aggressive than FS1022 (e.g. disease severity with FS1008 was twice that with FS1022 for inoculum concentrations of 5.10^3^ to 5.10^5^ conidia.mL^−1^). From inoculum concentrations of 5.10^3^ to 5.10^4^ conidia.mL^−1^, FS1008 induced a less severe attack if the other, less aggressive isolate (FS1022) was present at the same concentration. This difference was significant provided that the number of FS1022 conidia was the same. It disappeared at the highest inoculum concentrations (> 10^5^ conidia.mL^−1^, corresponding to a disease severity of 35-40%). Furthermore, the results for the mixture are actually very close to what would be seen with the same inoculum pressure (10^4^ and 10^5^ conidia.mL^−1^) with only the less aggressive isolate (FS1022).

**Fig. 8.**
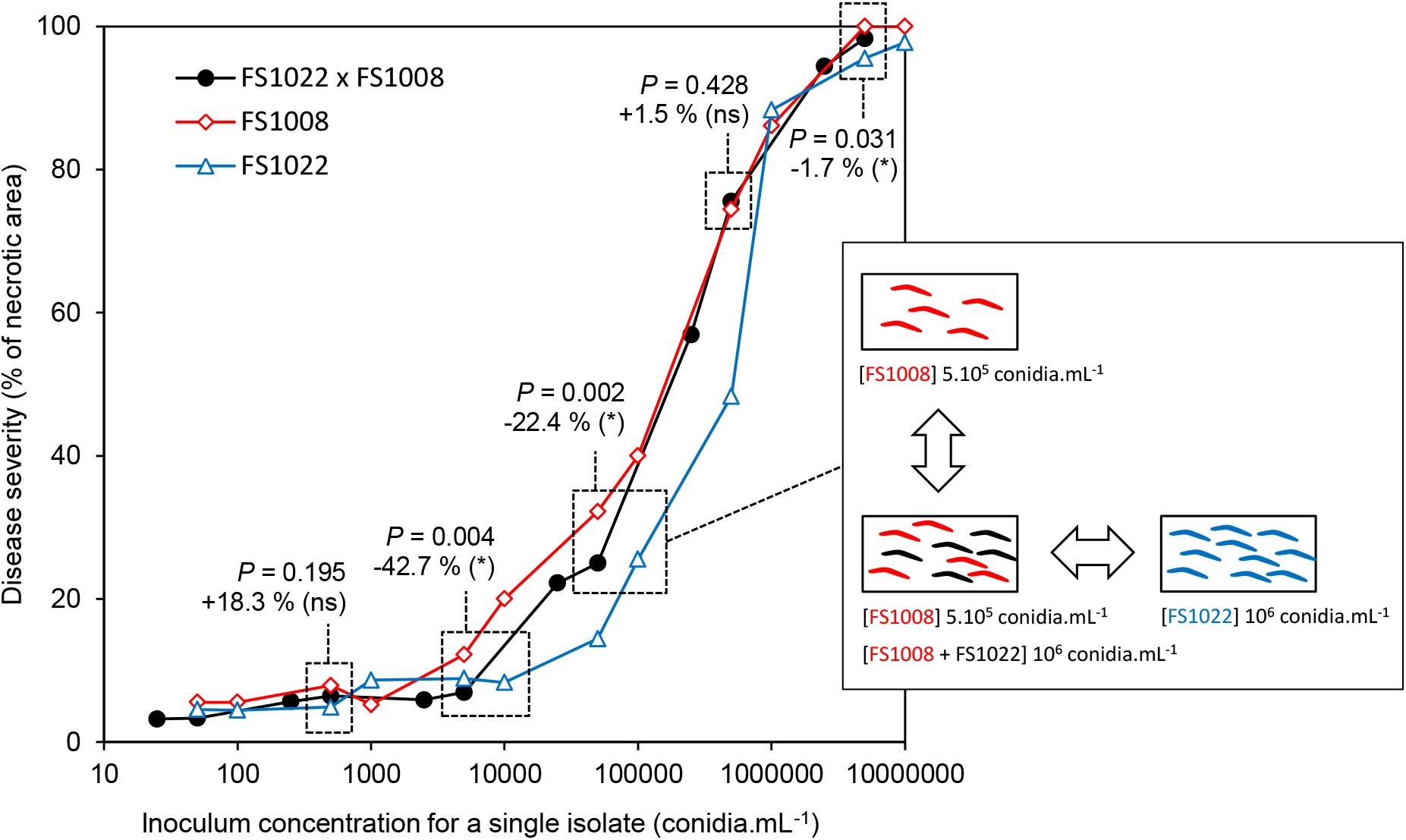
Relationship between inoculum concentration of *Zymoseptoria tritici* FS1008 or FS1022 isolate (number of FS1008 or FS1022 conidia per mL) in suspension alone or mixed with a second isolate present in the same amount, and the mean severity of Septoria tritici blotch (percentage of necrotic area) estimated on the three uppermost leaves of adult wheat plants cv. Soissons. ANOVA was performed (see *P*-value) at each relevant inoculum concentration (5.10^2^, 5.10^3^, 5.10^4^, 5.10^5^ and 5.10^6^ conidia of each isolate per mL; see example 5.10^5^ conidia of FS1008 in the scheme on the right). The number indicates the difference (in %) between the severity of the disease induced by FS1008 (the most aggressive isolate) inoculated alone and the severity of the disease induced by the FS1008 × FS1022 mixture (for the same number of FS1008 conidia); a negative percentage indicates that disease severity was lower after inoculation with the mixture; * indicates that the difference is significant (post hoc multiple comparison).

### Competition with asexual multiplication for resource allocation affects sexual reproduction in *Z. tritici*

Ascospores were collected after each of the five crosses in 2015, 2016 and 2017. The number of ascospores was largest for the FS802 × FS1006 cross in 2015 and lowest for the FS808 × FS806 cross in 2016 (Online Resource 2; Figure 9). Moreover, in 2016, the first ascospores were collected later in the season (January), due to relatively the dry weather from August to September 2016 (65 mm of rainfall, versus the mean of 170 mm generally observed this period). A similar effect of year was also observed with the ascomycetes *Leptosphaeria maculans* (Marie-Hélène Balesdent, INRA BIOGER, pers. comm.) and *Septoria linicola* (Annette Penaud, Terres Inovia, pers. comm.) in the same experimental area.

**Fig. 9.**
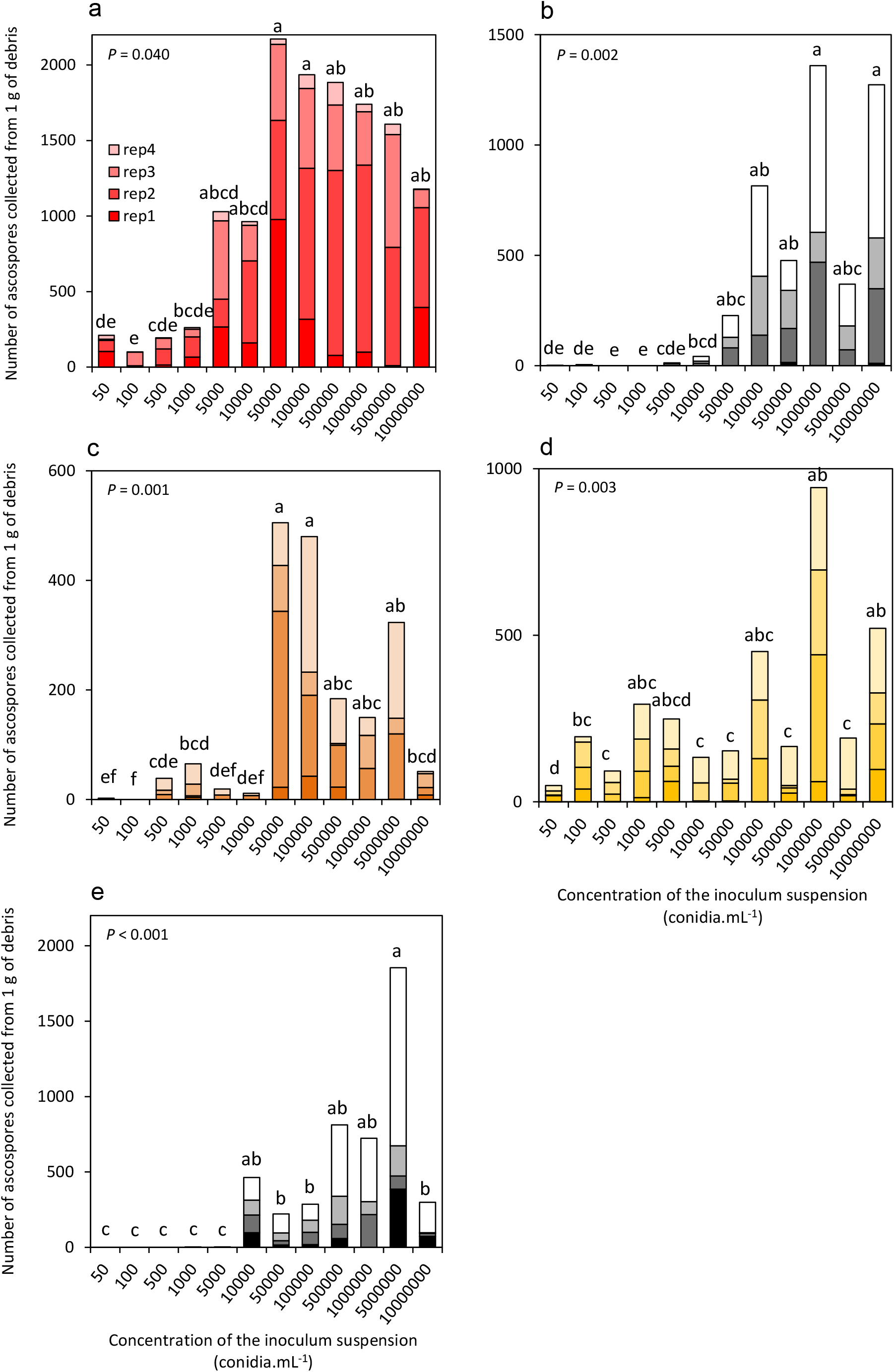
Impact of inoculum concentration (total number of conidia per mL) on the intensity of sexual reproduction in *Zymoseptoria tritici* (number of ascospores per g of wheat debris). Inoculation with biparental suspensions of isolates (a) FS802 × FS1006 (2015), (b) FS813 × FS732 (2016), (c) FS808 × FS806 (2016), and (d) FS1022 × FS1008 (2017) were performed on cv. Soissons, and with (e) 3D7 × 1A5 (2017) on cv. Runal. Each bar corresponds to the cumulative number of ascospores collected per gram of wheat debris over four independent discharge events (rep1 to rep4). For each cross, Kruskal-Wallis test was performed and completed, when significant (see P-value), by a post hoc multiple comparison; letter indicates significant differences between inoculum concentrations; tests for FS802 × FS1006 (2015) were performed after exclusion of rep4 data (to avoid bias caused by the overall low amount of ascospores). The complete dataset is given in Table S1.

For all crosses, the number of ascospores discharged increased with inoculum concentration. The smallest numbers of ascospores were collected for the lowest inoculum concentrations (50 or 10^2^ conidia.mL^−1^). The number of ascospores discharged at the highest inoculum level (10^7^ conidia.mL^−1^) was however systematically below the maximum number. For crosses on Soissons, this maximum was reached for inoculum concentrations of 5 × 10^4^ conidia.mL^−1^ (FS802 × FS1006 and FS813 × FS732) to 10^6^ conidia.mL^−1^ (FS813 × FS732 and FS1022 × FS1008). It was reached for an inoculum concentration of 5 × 10^6^ conidia.mL^−1^ for the cross performed on Runal (3D7 × 1A5), on which disease severity was significantly lower than on Soissons for inoculum levels of 5 × 10^2^ to 5 × 10^6^ conidia.mL^−1^ (Figure 5). This pattern was even more marked for the relationship between disease severity (proxy for asexual multiplication intensity) and the normalized number of ascospores (proxy for sexual reproduction intensity; Figure 10). The quadratic model and a model integrating two density-dependent constraints converged for three crosses (FS802 × FS1006, FS808 × FS806, FS1022 × FS1008), with the intensity of sexual reproduction increasing to a peak for disease severities of 35 to 50% and then decreasing thereafter. This pattern was particularly marked for the FS802 × FS1006 and FS808 × FS806 crosses. The initial ascending part of the curves illustrates an Allee effect (difficulty finding mates at low pathogen densities), whereas the second, descending part of the curve highlights competition for resource allocation between the two modes of reproduction (fewer host resources available for sexual reproduction due their previous mobilization for asexual multiplication).

**Fig. 10.**
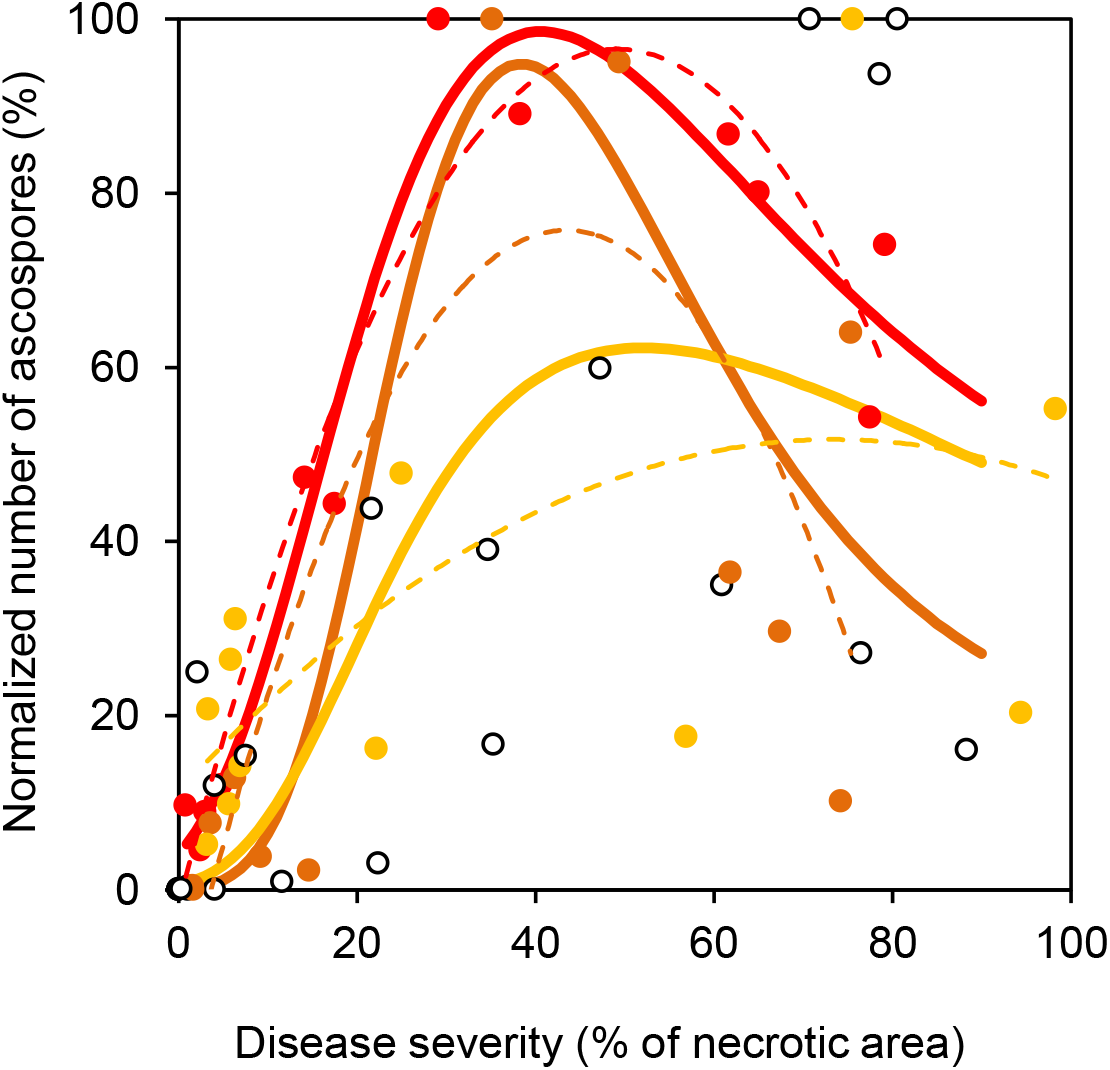
Empirical relationship between asexual multiplication intensity in *Zymoseptoria tritici* (disease severity assessed as a percentage of necrotic area) and sexual reproduction intensity (normalized number of ascospores for each cross expressed in %). A quadratic model (dotted lines; see eq. 2 in Material & Methods) and a model integrating two density-dependent constraints (bold lines; see eq. 3) were adjusted on the data for three crosses (FS802 × FS1006 in red, FS808 × FS806 in orange, FS1022 × FS1008 in yellow) to visualize the different relationships. Convergence was not obtained for the other two crosses (white points).

## Discussion

In field conditions, the different layers of wheat plants, from the soil to the head, contribute differently to *Z. tritici* reproduction. We showed that mean *Z. tritici* ascospore production was maximal between 25 and 35 cm above the ground. This significant impact of vertical positioning is complementary to the findings of Pfender & Wootke [27], who established that the pseudothecia of *Pyrenophora tritici-repentis* survived well in standing stubble and upper mulch layers, but not in straw or mulch located in the soil or directly on its surface. At least two antagonistic mechanisms are likely to account for the increase and subsequent decrease in sexual reproduction intensity with increasing height of the canopy layer above the ground. Higher positions in the canopy are associated with a lower density of lesions. The probability of crosses occurring, therefore, also decreases with height. Furthermore, higher positions in the canopy are occupied by younger wheat tissues, on which infections are likely to be more recent, with a lower probability of ascosporogenesis having had time to occur. Lower down in the canopy, the wheat tissues are older and the pseudothecia are already empty or degraded. These empirical observations thus complete the results obtained by reanalyzing the dataset of Eriksen & Munk [16].

Traits relating to asexual multiplication *in planta*, such as the density and exudation capacity of pycnidia, are affected by competition between the two parental isolates for resources. This result is consistent with the lower level of pycnidial coverage observed after inoculation with mixtures of isolates than after inoculation with single isolates [28, 29]. The competitive disequilibrium between the two parental isolates may also account for the density dependence of asexual development in *Z. tritici.* This finding is consistent with previous results obtained in the same experimental conditions in a previous study [23], in which an interval of two to three weeks between inoculation with the first parental isolate and inoculation with the second was found to be detrimental to ascosporogenesis. In this case, the host tissues were likely to be colonized by the first isolate, leaving fewer host resources available for the second isolate, consistent with the establishment of competition during the asexual stage of the disease cycle. Interactions of this type were studied by Clement et al. [8], who identified two basic response patterns, consisting of an increase or decrease in reproductive fitness in multiple infections of potato leaves with *P. infestans*, depending on pathogen genotype. In the case of *Z. tritici*, the activation of host defenses by the less virulent or less aggressive isolate may limit the subsequent development of a second more aggressive isolate at low inoculum pressure. Conversely, at high inoculum pressure, the host sites for infection are likely to be saturated, regardless of the aggressiveness of the isolates causing the infection. The less aggressive isolate (i.e. to which the host is less susceptible) may trigger defense responses that ultimately prevent the more aggressive isolate from developing to the extent that it would have if used alone for inoculation. The highest inoculum pressure was so high that this mechanism was not expressed.

Cirrhi produced by different isolates may contain different densities of pycnidiospores or may be produced later or earlier than five weeks after inoculation, and thus could bias the analysis. These differences between isolates may also be affected by environmental factors. Such aspects are beyond the scope of the current work but should be taken into account more accurately in further studies. Moreover, other reasons than competition for resources sensu stricto may explain the negative relationships between mean disease severity and the density of pycnidia or cirrhi within lesions. The presence of necrotic leaf tissue and the presence of pycnidia and cirrhi are the result of different developmental processes of the fungus and of different stages of the plant-fungus interaction [30]. Therefore, we can imagine a case where a proliferation of necrotic tissue does not lead to a proliferation of pycnidia due to the triggering of defence responses or to fungal auto-inhibition.

Sexual reproduction is driven by at least two density-dependent mechanisms with opposite impacts: a positive Allee effect due to mate limitation, and with a negative effect due to competition with asexual multiplication for resources. We provide here the first experimental evidence that sexual reproduction in a fungal foliar pathogen is driven by antagonistic density-dependent mechanisms.

*Positive density dependence of sexual reproduction* – Higher densities of *Z. tritici* lesions on leaves and stems are associated with a higher probability of encounter and a higher intensity of sexual reproduction. Mate limitation has been identified as the most common mechanism underlying Allee effects [4]. This experimental result is supported by the positive correlations previously established in natural conditions at both the field and landscape scales: ascospore production is generally greatest after the most severe epidemics [21, 22].

*Negative density dependence of sexual reproduction* – Strong Septoria tritici blotch attacks (i.e intense asexual multiplication) limit the resources available for sexual reproduction, resulting in a detrimental effect. We showed that competition between the two parental isolates during asexual multiplication is plausible and may lead to an imbalance between mating types. However, this explanation appears to be less relevant in natural conditions, given the high diversity of the pathogen population even at the leaf scale (sex ratio close to 1). As illustrated by the negative impact on fruiting and sporulating capacity, competition also limits the potential development of parental isolates in a similar way: host resources are overexploited during the asexual phase, to the detriment of sexual reproduction.

The optimal equilibrium between asexual and sexual reproduction in semi-controlled conditions corresponds to a disease severity of 30 to 45%. This optimum severity is lower than that observed in the field on the leaves of the lower plant layers, on which severity may reach very high levels (80-100%) and which senesce faster than the leaves of the upper layers.

Eriksen & Munk [16] reported that pseudothecia form only at disease severities exceeding 38%, following the coalescence of lesions, and that the highest proportion of pseudothecia occurs on leaves with a disease severity of 50 to 80%. Furthermore, the equation we fitted to these experimental data revealed no negative density-dependent effect, perhaps because the epidemiological survey stopped earlier (August) than in our own experiment conditions, suggesting that the pseudothecia may not have been mature in the upper plant layers.

As a final step in this study, we developed an overall synthetic representation of the relationship between the investments of *Z. tritici* in the two modes of reproduction at the host-tissue scale, taking into account the ecological processes discussed above. In Figure 11, we propose a theoretical relationship between asexual multiplication intensity (disease severity) and sexual reproduction intensity (ability to produce offspring). The number of fruitful encounters between parental isolates initially increases with increasing disease severity (1). During this first stage, competition between isolates may lead to mating-type disequilibrium, with an impact on sexual reproduction (2). At disease severities of more than 30-45%, a high-density pathogen population becomes disadvantageous, due to competition for host resources between the asexual and sexual modes of reproduction (3). This conceptual model is complementary to the findings of Suffert et al. [23], formalized by the title “Fashionably late partners have more fruitful encounters”. We can now extend the metaphor by providing *Z. tritici* with additional advice for even more fruitful encounters: “Do not live as a hermit but avoid crowded places!” This view was already put forward by Kokko & Rankin [1], in their study entitled “Lonely hearts or sex in the city?”

**Fig. 11.**
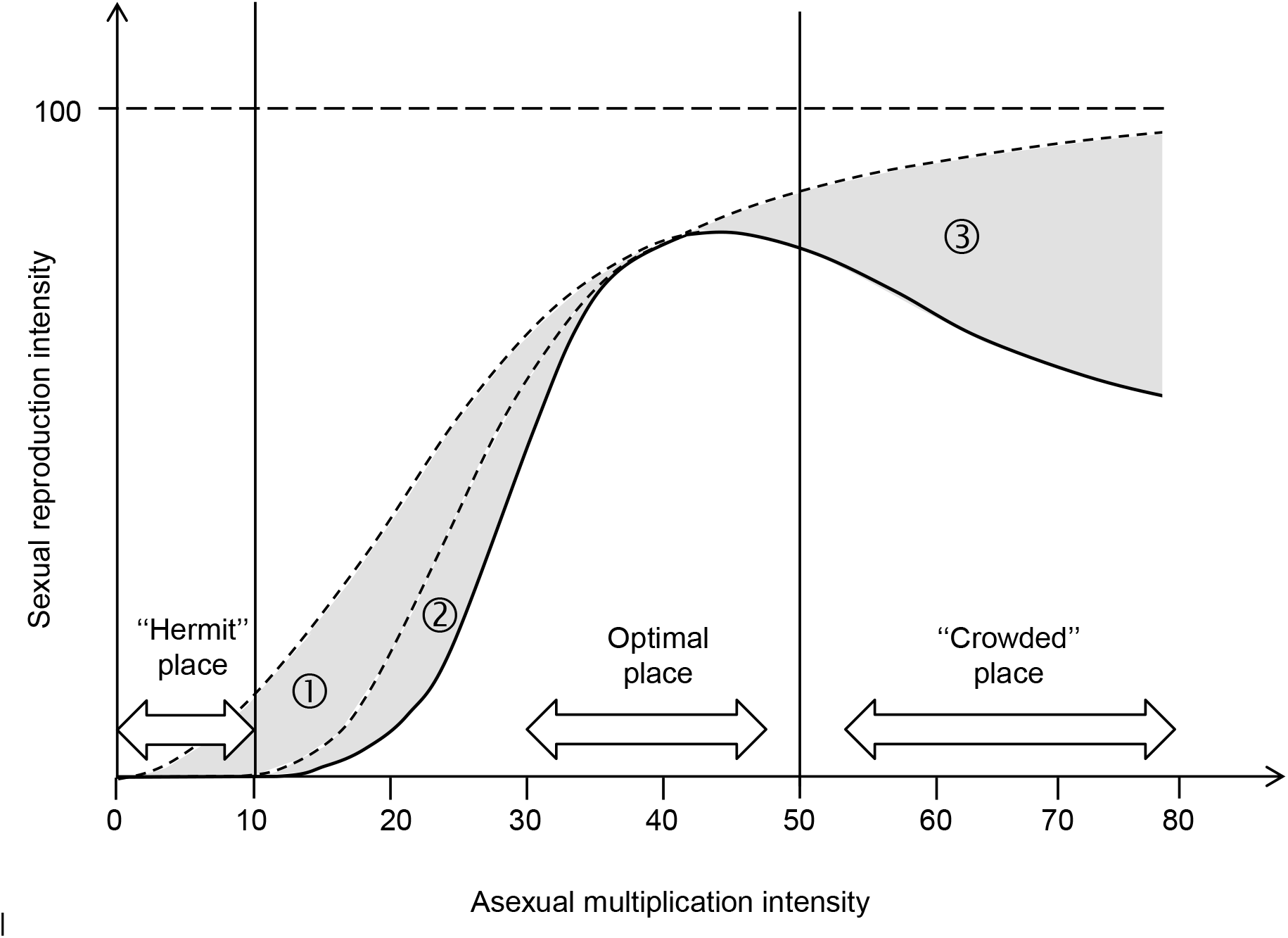
Theoretical relationship between asexual multiplication intensity in *Zymoseptoria tritici* (disease severity assessed as a percentage of necrotic area) and sexual reproduction intensity (normalized number of ascospores produced per g of wheat tissue), assuming that the conidia of two compatible parental isolates were deposited in equal proportions on leaves. Grey areas represent the decrease in sexual reproduction intensity due to: (1) an Allee effect (difficulty finding the opposite mating type at low population densities); (2) induced resistance (activation of host defense mechanisms by one of the parental isolates, leading to a mating-type disequilibrium); (3) competition for host resource allocation between asexual multiplication and sexual reproduction. Sexual reproduction intensity is expected to be optimal for disease severities of 30 to 45%.

Is an optimal disease severity of 30-45% an appropriate ecological equilibrium determining the evolutionary endpoints of selection in *Z. tritici*? This theoretical question needs to be addressed in an evolutionary ecology perspective, and we can now use empirical data to try to answer it. Indeed, a successful parasitic life depends on optimal exploitation of the host to satisfy key functions directly involved in reproductive fitness. Theoretically, both weakly aggressive isolates causing only a few small lesions, and very aggressive isolates causing large numbers of large lesions, have a lower probability of transmitting a part of their genetic background by sexual reproduction than isolates with intermediate levels of aggressiveness. This view is complementary to the findings of Kema et al. [19], which explained the maintenance of avirulent individuals in the pathogen population despite the widespread deployment of resistance genes (e.g. AvrStb6 despite the extensive presence of Stb6 in the wheat lines used in breeding programs worldwide [31]; Thierry Marcel, INRA BIOGER, com. pers.). Sexual reproduction therefore plays a key role in both the evolution of virulence and aggressiveness.

A previous study [23] revealed an absence of functional trade-offs between the two modes of reproduction in *Z. tritici:* no adaptive compromise was established between pathogenicity and transmission in analyses of pathogen life traits at the individual level. An overall epidemiological trade-off was, however, established between intra- and interannual scales over a larger spatiotemporal scale (field and surrounding area), probably driven by the consequences of sexual reproduction for local population dynamics relating to selection and counter-selection [22]. However it is not clear that the optimal equilibrium between sexual and asexual reproduction corresponds to the maximal intensity of sexual reproduction, as one could think; asexual reproduction may be for instance more beneficial to overall strain fitness in the early stages of an epidemic, where inoculum (estimated for instance by the number of infection units available per m^2^ of crop) is more limited than during the spring period [32]. According to the findings reported here, the potential antagonism between the asexual and sexual modes of reproduction and its epidemiological consequences appear to be more complex, due to a dependence on scale, from host tissues up to agricultural landscape scale.

Sexual reproduction is clearly a key process in the eco-epidemiology of Septoria tritici blotch. An understanding of its determinants may open up new perspectives for the management of other foliar fungal pathogens with dual modes of reproduction.

---

**Online Resource 2.** Number of *Zymoseptoria tritici* ascospores collected per gram of wheat debris during four independent discharge events for the five crosses (FS802 × FS1006, FS813 × FS732, FS808 × FS806, FS1022 × FS1008, 3D7 × 1A5) performed with 12 inoculum concentrations.

| FS802 × FS1006 on wheat cv. Soissons (2015) |  |  |  |  |  |  |  |
| --- | --- | --- | --- | --- | --- | --- | --- |
| Inoculum concentration <sup>1</sup> | Disease severity <sup>2</sup> | Mean number of ascospores collected per gram of debris |  |  |  |  |  |
| Date | 05-02-2015 | 11-12-2015 | 11-19-2015 | 12-03-2015 | 01-21-2016 | - | Total |
| 50 | 0.8 ± 0.3 | 103.2 | 72.5 | 7.9 | 26.6 |  | 210.2 |
| 10<sup>2</sup> | 2.5 ± 0.6 | 0.3 | 7.5 | 91.2 | 0.3 |  | 99.2 |
| 5 × 10<sup>2</sup> | 3.0 ± 0.5 | 13.5 | 105.6 | 71.5 | 3.7 |  | 194.3 |
| 10<sup>3</sup> | 4.8 ± 1.3 | 66.4 | 132.9 | 51.2 | 11.4 |  | 261.9 |
| 5 × 10<sup>3</sup> | 14.2 ± 2.2 | 265.0 | 184.6 | 518.7 | 60.9 |  | 1029.1 |
| 10<sup>4</sup> | 17.5 ± 3.5 | 159.2 | 543.2 | 236.3 | 24.6 |  | 963.3 |
| 5 × 10<sup>4</sup> | 29.2 ± 3.4 | 976.8 | 656.1 | 504.4 | 35.5 |  | 2172.7 |
| 10<sup>5</sup> | 38.3 ± 4.7 | 316.2 | 999.9 | 529.3 | 90.8 |  | 1936.1 |
| 5 × 10<sup>5</sup> | 61.7 ± 3.1 | 76.3 | 1225.2 | 433.8 | 150.0 |  | 1885.2 |
| 10<sup>6</sup> | 65.0 ± 4.6 | 98.7 | 1238.5 | 353.6 | 49.9 |  | 1740.7 |
| 5 × 10<sup>6</sup> | 79.2 ± 4.2 | 10.1 | 781.8 | 747.6 | 69.3 |  | 1608.8 |
| 10<sup>7</sup> | 77.5 ± 4.9 | 395.0 | 660.1 | 120.6 | 3.1 |  | 1178.7 |

<sup>1</sup> total number of conidia per mL
<sup>2</sup> percentage of necrotic area ± standard deviation

FS813 × FS732 on wheat cv. Soissons (2016)
| Inoculum<br>concentration <sup>1</sup> | Disease<br>severity <sup>2</sup> | Mean number of ascospores collected per gram of debris |  |  |  |  |  |
| --- | --- | --- | --- | --- | --- | --- | --- |
| Date | 05-06-2016 | 11-17-2016 | 01-19-2017 | 02-15-2017 | 03-01-2017 | 03-15-2017 | Total |
| 50 | 1.1 ± 0.4 | 0.0 | 0.0 | 2.0 | 0.0 | 0.0 | 2.0 |
| 10<sup>2</sup> | 1.0 ± 0.5 | 0.0 | 0.0 | 0.0 | 4.5 | 0.0 | 4.5 |
| 5 × 10<sup>2</sup> | 0.9 ± 0.3 | 0.0 | 0.0 | 0.0 | 0.0 | 0.0 | 0.0 |
| 10<sup>3</sup> | 1.6 ± 0.4 | 0.0 | 0.0 | 0.0 | 0.0 | 0.0 | 0.0 |
| 5 × 10<sup>3</sup> | 9.3 ± 2.0 | 0.0 | 0.0 | 9.6 | 0.2 | 2.9 | 12.7 |
| 10<sup>4</sup> | 17.1 ± 2.6 | 0.0 | 0.0 | 10.6 | 10.0 | 21.2 | 41.8 |
| 5 × 10<sup>4</sup> | 33.3 ± 2.1 | 0.0 | 0.0 | 80.6 | 47.8 | 98.5 | 227.0 |
| 10<sup>5</sup> | 41.3 ± 3.8 | 0.0 | 0.0 | 138.8 | 266.9 | 409.0 | 814.7 |
| 5 × 10<sup>5</sup> | 57.2 ± 3.7 | 0.0 | 14.4 | 154.7 | 172.4 | 134.8 | 476.3 |
| 10<sup>6</sup> | 67.8 ± 1.7 | 0.0 | 0.7 | 467.9 | 135.4 | 755.9 | 1360.0 |
| 5 × 10<sup>6</sup> | 75.6 ± 1.5 | 0.0 | 0.0 | 72.2 | 108.1 | 189.6 | 369.9 |
| 10<sup>7</sup> | 78.3 ± 2.4 | 0.0 | 10.6 | 338.2 | 230.3 | 694.5 | 1273.7 |

<sup>1</sup> total number of conidia per mL
<sup>2</sup> percentage of necrotic area ± standard deviation

FS808 × FS806 on wheat cv. Soissons (2016)
| Inoculum<br>concentration <sup>1</sup> | Disease<br>severity <sup>2</sup> | Mean number of ascospores collected per gram of debris |  |  |  |  |  |
| --- | --- | --- | --- | --- | --- | --- | --- |
| Date | 05-06-2016 | 11-17-2016 | 01-19-2017 | 02-15-2017 | 03-01-2017 | 03-15-2017 | Total |
| 50 | 1.6 ± 0.3 | 0.0 | 0.0 | 0.2 | 2.1 | 0.0 | 2.3 |
| 10<sup>2</sup> | 1.4 ± 0.3 | 0.0 | 0.0 | 0.0 | 0.0 | 0.0 | 0.0 |
| 5 × 10<sup>2</sup> | 2.8 ± 0.7 | 0.0 | 0.0 | 9.0 | 7.6 | 21.9 | 38.5 |
| 10<sup>3</sup> | 5.9 ± 1.5 | 0.0 | 4.1 | 2.6 | 21.5 | 36.9 | 65.0 |
| 5 × 10<sup>3</sup> | 7.3 ± 1.5 | 0.0 | 0.0 | 8.1 | 0.0 | 11.1 | 19.2 |
| 10<sup>4</sup> | 9.8 ± 2.6 | 0.0 | 0.0 | 6.9 | 0.0 | 4.2 | 11.2 |
| 5 × 10<sup>4</sup> | 28.9 ± 2.6 | 0.0 | 22.2 | 321.2 | 83.9 | 78.0 | 505.3 |
| 10<sup>5</sup> | 43.9 ± 3.1 | 0.0 | 42.6 | 147.7 | 42.3 | 247.6 | 480.2 |
| 5 × 10<sup>5</sup> | 60.0 ± 2.9 | 0.0 | 22.7 | 76.5 | 3.1 | 81.8 | 184.0 |
| 10<sup>6</sup> | 65.0 ± 3.3 | 0.0 | 0.0 | 56.7 | 60.2 | 32.8 | 149.7 |
| 5 × 10<sup>6</sup> | 75.6 ± 1.6 | 0.0 | 0.6 | 119.2 | 28.3 | 175.4 | 323.5 |
| 10<sup>7</sup> | 74.4 ± 2.2 | 0.0 | 8.2 | 13.6 | 25.5 | 4.0 | 51.2 |

| Inoculum<br>concentration <sup>1</sup> | Disease<br>severity <sup>2</sup> | Mean number of ascospores collected per gram of debris |  |  |  |  |  |
| --- | --- | --- | --- | --- | --- | --- | --- |
| Date | 05-27-2017 | 11-09-2017 | 12-07-2017 | 01-04-2017 | 01-01-2018 | 01-24-2018 | Total |
| 50 | 3.2 ± 0.2 | 0.0 | 18.3 | 1.7 | 12.7 | 16.1 | 48.8 |
| 10<sup>2</sup> | 3.3 ± 0.2 | 0.0 | 38.8 | 64.6 | 75.6 | 16.8 | 195.7 |
| 5 × 10<sup>2</sup> | 5.7 ± 0.5 | 0.0 | 0.9 | 22.0 | 35.0 | 34.8 | 92.8 |
| 10<sup>3</sup> | 6.4 ± 0.7 | 0.0 | 12.5 | 79.2 | 96.6 | 104.8 | 293.1 |
| 5 × 10<sup>3</sup> | 5.9 ± 0.5 | 0.0 | 61.2 | 45.5 | 51.4 | 90.9 | 249.0 |
| 10<sup>4</sup> | 6.9 ± 1.2 | 0.0 | 2.2 | 0.4 | 54.2 | 77.0 | 133.8 |
| 5 × 10<sup>4</sup> | 22.2 ± 2.2 | 0.0 | 2.8 | 53.6 | 11.6 | 84.9 | 152.8 |
| 10<sup>5</sup> | 25.0 ± 2.4 | 0.0 | 0.1 | 129.5 | 176.1 | 145.6 | 451.3 |
| 5 × 10<sup>5</sup> | 56.9 ± 3.6 | 0.0 | 26.2 | 15.5 | 7.3 | 117.0 | 166.0 |
| 10<sup>6</sup> | 75.6 ± 5.0 | 0.0 | 60.6 | 381.1 | 254.4 | 247.5 | 943.6 |
| 5 × 10<sup>6</sup> | 94.4 ± 0.4 | 0.0 | 18.4 | 3.2 | 16.3 | 153.5 | 191.4 |
| 10<sup>7</sup> | 98.3 | 0.0 | 97.3 | 136.4 | 92.9 | 194.3 | 521.0 |

<sup>1</sup> total number of conidia per mL
<sup>2</sup> percentage of necrotic area ± standard deviation

3D7 × 1A5 on wheat cv. Runal (2017)
| Inoculum<br>concentration <sup>1</sup> | Disease<br>severity <sup>2</sup> | Number of ascospores collected per gram of debris |  |  |  |  |  |
| --- | --- | --- | --- | --- | --- | --- | --- |
| Date | 05-27-2017 | 11-09-2017 | 12-07-2017 | 01-04-2017 | 01-01-2018 | 01-24-2018 | Total |
| 50 | 0.0 | 0.0 | 0.0 | 0.0 | 0.0 | 0.0 | 0.0 |
| 10<sup>2</sup> | 0.0 | 0.0 | 0.0 | 0.0 | 0.0 | 0.0 | 0.0 |
| 5 × 10<sup>2</sup> | 0.0 | 0.0 | 0.0 | 0.0 | 0.0 | 0.0 | 0.0 |
| 10<sup>3</sup> | 0.0 | 0.0 | 0.0 | 0.0 | 0.0 | 2.2 | 2.2 |
| 5 × 10<sup>3</sup> | 0.3 ± 0.1 | 0.0 | 0.0 | 1.4 | 0.0 | 0.2 | 1.6 |
| 10<sup>4</sup> | 2.1 ± 0.3 | 0.0 | 97.6 | 117.2 | 98.0 | 150.2 | 463.0 |
| 5 × 10<sup>4</sup> | 4.1 ± 0.2 | 0.0 | 15.5 | 29.5 | 51.4 | 124.8 | 221.2 |
| 10<sup>5</sup> | 7.6 ± 0.3 | 0.0 | 19.2 | 81.0 | 79.5 | 106.2 | 286.0 |
| 5 × 10<sup>5</sup> | 21.7 ± 0.7 | 0.0 | 58.4 | 95.0 | 185.0 | 473.8 | 812.1 |
| 10<sup>6</sup> | 34.7 ± 2.8 | 0.0 | 0.0 | 217.4 | 85.8 | 420.5 | 723.7 |
| 5 × 10<sup>6</sup> | 80.6 ± 0.4 | 0.0 | 386.2 | 87.5 | 200.2 | 1181.4 | 1855.3 |
| 10<sup>7</sup> | 88.3 ± 1.8 | 0.0 | 73.2 | 19.6 | 4.2 | 201.5 | 298.5 |

<sup>1</sup> total number of conidia per mL
<sup>2</sup> percentage of necrotic area ± standard deviation

**Online Resource 2.**
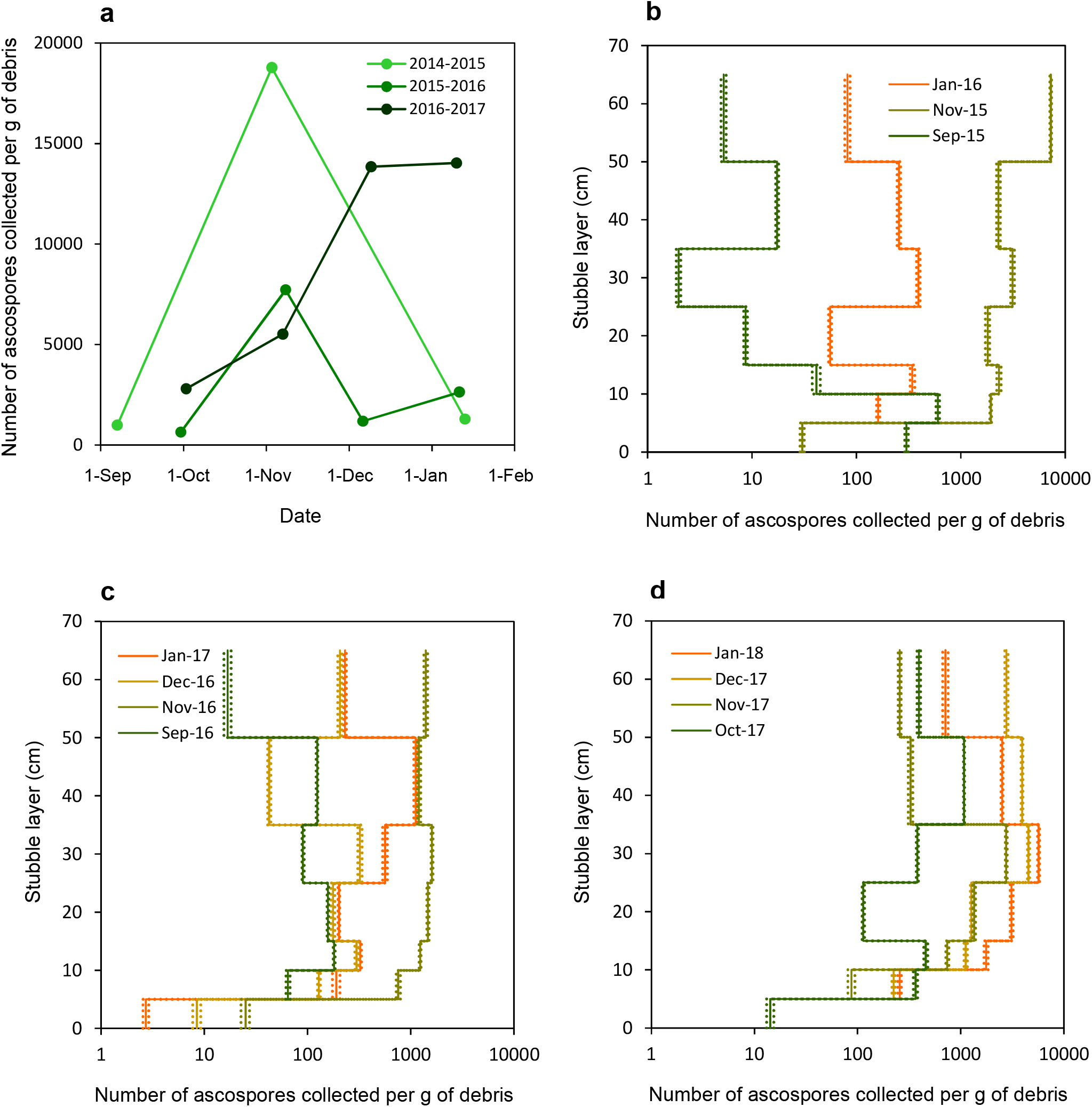
(a) Spatiotemporal dynamics of *Zymoseptoria tritici* sexual reproduction in standing stubble left in the field during the fall, over three interepidemic periods (2015–2016, 2016-2017, 2017-2018). (b, c, d) Impact of the vertical position of wheat tissues (stems and leaves, from lower to upper layers: 0-5 cm, 5-10 cm, 10-15 cm, 15-25 cm, 25-35 cm, 35-50 cm, 50-65 cm) on the number of ascospores collected per gram of debris (logarithmic scale) on three or four sampling dates (from September to January) over the three interepidemic periods. The dotted lines indicate the standard error.

## Acknowledgments

We thank Nathalie Retout (INRA BIOGER, France) for technical assistance, Dr. Frédéric Hamelin (Agrocampus Ouest, France) for preliminary discussions on ecological modeling aspects, and Anne-Lise Boixel (INRA BIOGER, France) for her help in statistical analyses. We thank Julie Sappa for her help correcting our English. We thank Dr. Alexey Mikaberidze and Prof. Bruce McDonald (ETH Zürich, Switzerland) for providing the ST99CH-3D7 and ST99CH-1A5 strains of *Z. tritici* and seeds of the wheat cv. Runal. We thank the two anonymous reviewers for their constructive comments, which helped us to improve the manuscript. This study was supported by a grant from the European Union Horizon Framework 2020 Program (Grant Agreement no. 634179, EMPHASIS project) covering the 2015-2019 period.

